# Active Transcriptome Remodeling and Epigenetic Adaptations Underlie Persistence in *Leishmania* parasites

**DOI:** 10.64898/2026.05.29.726936

**Authors:** Eyson Quiceno, Cristian Camilo Rodríguez-Almonacid, Khalid Omeir, Jacob Ancira, Caleb Phillips, Zemfira N. Karamysheva

## Abstract

Persistence - characterized by the transient ability of subpopulations of drug-susceptible parasites to survive exposure to drug - is a major driver of treatment failure and clinical relapses in leishmaniasis. Persisters are characterized by non-dividing or slow-growing state and increased drug tolerance. However, the molecular mechanisms governing formation of persisters remain poorly understood in *Leishmania* parasites. Here, we developed a model to explore persistence in *Leishmania mexicana*. Viable promastigote persister-like subpopulations were enriched using Ficoll density gradient centrifugation following lethal exposure to antimonial drugs that killed 80% of parasites. The surviving parasites exhibited delayed growth in drug-free medium that is characteristic for persisters, and significantly higher tolerance to drug upon rechallenge. Transcriptomic profiling across acute stress, drug-free recovery, and rechallenge phases revealed a global remodeling in persisters under all tested conditions. Induction phase in *Leishmania* persisters was characterized by downregulation of several biological processes and a robust upregulation of nucleolar pathways, supporting epitranscriptomic changes during formation of persisters. Upon drug removal, this profile rapidly reverted, initiating ribosomal biogenesis to exit latency and resume cellular proliferation. Resuscitation phase exhibited activated protein synthesis and upregulation in many biological processes associated with metabolic and mitochondrial functions. Furthermore, comparative analysis of drug responses in rechallenged drug-tolerant persisters and parental parasites exposed to the drug for the first time, revealed that persisters exhibit distinct drug response profiles compared to parental parasites by rapidly implementing a highly conserved, coordinated survival reprogramming, where 316 genes were uniquely downregulated, and 241 genes were upregulated. The distinct features of the drug response in rechallenged persister cells were characterized by the downregulation of mitochondrial function and protein synthesis machinery to induce a dormant, idling state, and the upregulation of drug-response and stress-tolerance genes to survive immediate toxicity. In contrast, parental parasites displayed a broad and disorganized drug response. Additionally, rechallenged persisters exhibited a distinct transcriptomic memory that transiently phenocopies stable genetic resistance. This pre-adapted state is characterized by the targeted upregulation of epigenetic modulators, heavy metal transporters, and catabolic enzymes to maintain viability. These findings demonstrate that drug persistence in *Leishmania* is not merely a metabolic collapse, but rather a sophisticated survival strategy involving active transcriptome remodeling, downregulation of translation and epigenetic adaptations. This transient state constitutes an initial evolutionary step toward permanent drug resistance and highlights new molecular vulnerabilities for therapeutic interventions aimed at preventing clinical relapse.

**Author summary:** The efficacy of leishmaniasis chemotherapy is often fails because *Leishmania* parasites survive the drugs by entering a dormant state, causing relapses in patients. In this study, however, we discovered that these parasites are not simply ‘quiescent’. On the contrary, they orchestrate a highly active survival response: they shut down energy-consuming functions while keeping their cellular alert mechanisms active. Remarkably, upon re-exposure to antileishmanial drugs, the parasites recall previous stress events and transiently activate defensive pathways characteristic of stable drug resistance. Understanding how *Leishmania* employs this temporary adaptive strategy provides critical targets for developing therapies designed to eradicate the infection and avert relapses.

## Introduction

*Leishmania* species cause leishmaniasis, a major health problem that affect around 12 million people worldwide (1) and is present in more than 98 countries (2). The incidence of leishmaniasis is on the rise in U.S. due to population growth, climate change, and increased travel. While there is consensus that leishmaniasis is a uniquely tropical disease, evidence from recent studies indicate that 59% of cases are endemic in southern states including Texas, where it is prevalent and has a genotypically distinct *Leishmania mexicana* strain (3–5). Control of leishmaniasis is hampered by the absence of a safe vaccine, inherent limitations of current medications, drug toxicity, and frequent treatment failures stemming from drug resistance and the pathogen’s persistence.

In general, persistence is defined as an adaptive phenotype where the organism exhibits a transient increase in tolerance to different stress conditions such as drugs, reactive oxygen species (ROS), pH, temperature, starvation or interaction with the host immune system. Persisters are known to be present in bacteria, fungal pathogens and protozoa parasites. However, persister cells can also be found in cancer, leading to cancer relapse and drug resistance (6). During this state the cell typically adopts a non-growing or slow growing state with reduced metabolism (7, 8); nevertheless, this phenotype is complex and heterogeneous, recent reports indicate that distinct subpopulations within persisters communities may retain the ability to replicate actively (9). Persisters are capable to exit a non-dividing state and return to the proliferative state when an insult, such as drug, is gone. Therefore, drug persistence in pathogens and cancer cells represents a major clinical challenge. It contributes to treatment failure, drives the emergence of drug-resistant populations, and represents a serious global health threat (10).

While resistance arises from genomic mutations that enable cells to survive and proliferate in the presence of drugs (11), persistence is a non-heritable phenotypic variation. Persister cells remain genetically identical to non-tolerant cells (8, 12), and once the stressor is removed from the medium, those cells can resume their proliferation after certain time and exhibit sensitivity to the original stressor (7). For this reason, compared to resistance, persistence is considered a transient and reversible stage (13).

The mechanisms associated with generation of persistence are poorly understood. There is an ongoing debate regarding whether cell dormancy and low metabolism are merely consequences of environmental stressors or if they are regulated mechanisms that actively drive the population into persister state. In bacteria, the dormant state has been associated with the activation of stress pathways, allowing dormant cells to survive because the drug cannot fulfill its function (14). Furthermore, several specific mechanisms have been linked to bacterial persistence, including toxin-antitoxin (TA) modules, genes associated with metabolic regulation, inactivation of pathways targeted by antibiotics, proteins involved in heat shock response, trans-translation, the SOS response, the stringent response mediated by (p)ppGpp, efflux pumps, epigenetic modifications, RNA degradation, and small non-coding RNA modulating translation (7, 14, 15). Recently, deficiency in translation has been identified as one of the important mechanisms leading to dormancy and persistence in bacteria (16, 17). Cancer persisters mirror the bacterial model, emphasizing non-genetic mechanisms and a state of reduced proliferation. The persister state in cancer cells is highly plastic and reversible, often involving epigenetic modifications and metabolic shifts that enhance survival under drug pressure (18). These cells can exit the dormant state and resume proliferation once the drug pressure is removed, leading to tumor relapse. Cancer persisters don’t just stop protein production, they specifically boost the translation of mRNAs that promote survival and resistance (19). Thus, in both bacterial and cancer persisters, the role of translation involves a global suppression of protein synthesis combined with the selective translation of specific pro-survival mRNAs to adapt to stress and evade treatments. This selective control allows cells to enter a dormant or slow-cycling state, rather than dying, and eventually re-emerge to cause chronic infections or cancer relapse.

Persisters have been reported in protozoan pathogens such as *Plasmodium* spp (20, 21), *Trypanosoma* spp (22, 23), *Toxoplasma* spp, and *Leishmania* spp (11, 12, 24). This behavior is fundamental for parasites to establish persistent infections within different hosts and survive the environmental stress. For instance, during *Leishmania* infection in mammalian hosts, a population of persister parasites has been identified (25), represented as a small number of parasites that includes both rapidly replicating and quiescent subpopulations, where some cells exhibit active DNA replication, while others are largely dormant or slow dividing. These parasites exhibit remarkable resistance to hostile conditions, such as oxidative stress from nitric oxide (NO) generated by host immune cells, which may also confer high tolerance to antileishmanial drugs.

While the mechanisms associated with drug resistance in *Leishmania*, have been widely studied, ranging from a genetic component with mutations in key genes, to efflux transporters, metabolic adaptations, and translational control (26), the role and underlying causes of persistence in this parasite remain unclear. This represents a significant gap in current research. To address this, the present study established a model to study drug persistence in promastigotes of *Leishmania mexicana* and examined transcriptomic changes associated with persistence.

## RESULTS

### Experimental design to generate persister-like parasites

The study of persisters in *Leishmania* is hampered by the lack of suitable models and difficulties in their isolation. To address this, we have established conditions to generate persister-like parasites that enter a non-dividing or slow-growing state and exhibit enhanced drug tolerance. In summary, promastigote persisters were induced *in vitro* by using antimony drug at a concentration that killed 80% of parasites; surviving cells (persisters) were purified on Ficoll gradient, resuspended in drug-free medium for recovery (12h) and examined for cell proliferation in drug-free medium and drug tolerance upon rechallenge. Our experimental design comprised four stages: a) parasite culturing, b) drug challenge and isolation of survived parasites on Ficoll gradient, c) recovery in drug-free medium and d) re-challenge with antimony drug (Fig. 1).

**Figure 1.**
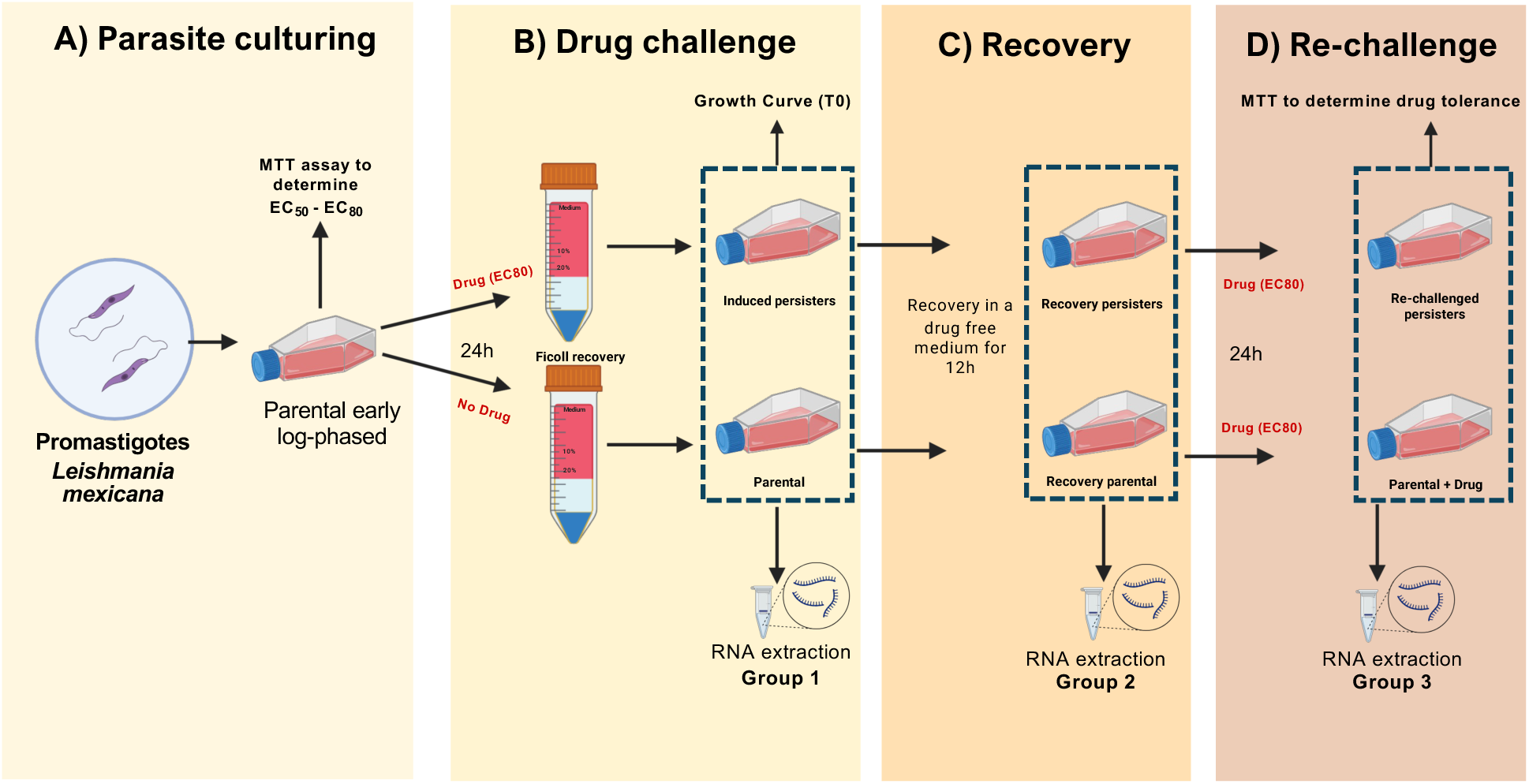
Experimental design for generating and evaluating *Leishmania mexicana* persister cells. The workflow comprises four stages (A-D): **(A)** Parasite culturing: *L. mexicana* promastigotes were seeded at 5×10^5^ cells/mL and collected at the early log phase to ensure high parasite viability (> 95%). **(B)** Drug challenge: the collected parasites were split into two groups – 1) parental parasites or the control group, which was never treated with the drug, and 2) treated group, a parasite population that was challenged with the drug (EC_80_ dose: 0.315 mM PAT) to kill 80% of parasites. Parasites that survived the treatment considered to be persisters. Both parental and treated live persisters were separated from dead cells on Ficoll gradient to reduce technical variability. **(C)** Recovery: after Ficoll isolation live persisters were twice washed to remove the drug remnant and recovered in a medium depleted of the drug. Similar treatment was applied to the control group. **(D)** Re-challenge: after recovery persisters were re-exposed to a second challenge with varying PAT doses (2, 1, 0.5, 0.25, 0.125, 0 mM). Parental group was subjected to the same drug challenge first time as a control. The colorimetric MTT cell viability assay was used to quantify improved tolerance compared to the parental control. The estimated time required per stage is described in hours. Trivalent antimony was administered as potassium antimonyl tartrate (PAT). Cells were collected for RNA extraction from groups 1-3 for RNA-seq and subsequent transcriptomic studies.

Initially, an MTT assay was performed on the parental strain exposed to varying PAT concentrations (0.125-2 mM) to determine the half-maximal effective concentration (EC_50_), defined as the drug concentration that reduces parasite viability by 50% (Fig. 1A and S1 Fig). Based on the parental strain’s non-linear regression model (EC50 = 0.1638 mM, Hill slope = -2.114; S1 Fig), we calculated an EC_80_ of 0.315 mM. This concentration, which reduces the live parasite population to 20% (persisters), was selected for all follow-up experiments.

### Persister-like parasites exhibit lower proliferation rate and significantly higher tolerance to drug

After establishing the EC_80_, we cultured susceptible *L. mexicana* promastigotes for 48 hours before splitting them into two conditions. The “Induced persisters” group received 0.315 mM PAT (EC_80_) for 24 hours, while the control group (“Parental”) remained untreated. After 24 hours of drug challenge, treated parasites displayed two different morphologies: apoptotic shape, characterized by rounded parasites; and regular slender morphology in live survivors, considered as persisters (S2 Fig). We isolated viable parasites (Induced persisters and Parental control; Fig. 1B) via Ficoll enrichment, obtaining aliquots for RNA extraction and growth curve assay (designated as T0 – time 0) (Fig. 1B). The growth rate was monitored in drug-free medium by over 96 hours, with follow-up measurements every 12 hours (Fig. 2A). Control samples reached a concentration peak of ∼5.1×10^7^ parasites/mL at hour 60, while treated samples showed a delayed proliferation reaching the peak of ∼3.5×10^7^ parasites/mL at hour 60 (Fig. 2A). Moreover, treated parasites couldn’t reach the same density as parental cells, supporting the existence of cells in dormant-like state. Parasites were then washed and allowed to recover in drug-free medium for 12 hours (Fig. 1C). After this time, we collected cell aliquots and performed RNA extraction on “Recovery persisters” and “Recovery parental” groups. Finally, we treated the “Recovery persisters” and matching parental control (Recovery Parental) with 0.315 mM PAT (EC_80_) and performed RNA extractions on “Rechallenged persisters” and “Parental + Drug” groups followed by RNA-seq for all acquired samples.

**Figure 2.**
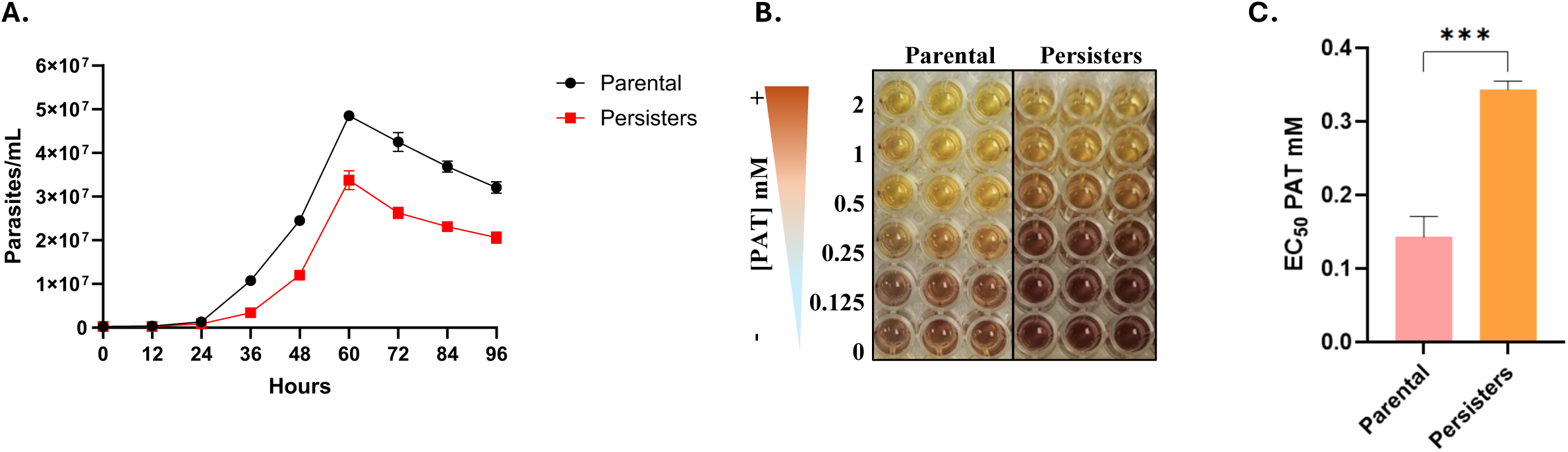
*Leishmania mexicana* persister-like promastigotes exhibit a lower proliferation rate and significantly higher drug tolerance. **(A)** The parasite proliferation was monitored during a recovery period of 96 hours by counting every 12 hours. Curves of growth show that persister parasites (red line) grow slower than the parental parasites (black line). Three independent biological replicates were measured. **(B)** The colometric MTT viability assay was used to measure the EC50 values for parental and persister parasites challenged with different drug concentrations in three biological replicates. Negative (-) or positive (+) symbols correspond to the presence/absence of drug. A representative experiment is shown. **(C)** EC_50_ values were calculated in three biological replicates using Graph prism and a Two-tailed T-test statistical method. Mean and standard error are plotted. The statistical difference was detected after comparing the EC_50_ values calculated for parental and persister parasites. value p <0.0001. The raw data are available in Source Data 1 (S1 Data).

At the same time, we evaluated drug tolerance by exposing these recovered populations to 0.125–2 mM PAT to calculate new EC50/EC80 values in “Rechallenged Persisters” (Fig. 1D). Drug tolerance in parental and persisters-like parasites was measured using MTT viability assay (Figs 1D, 2B-C and S3). The results revealed a significant increase in drug tolerance among the persisters-like parasites, demonstrating roughly two-fold difference between the persister and parental groups (Fig. 2B). Our findings indicate that persisters-like parasites exhibit typical characteristics of persistence: enhanced tolerance and a quiescent state, followed by gradual resumption of growth.

### Persister parasites exhibit extensive transcriptome remodelling

Three groups of RNA samples corresponding to induced persisters and parental control (group 1); recovery persisters and recovery parental control (group 2) and rechallenged persisters and parental + drug control exposed to the PAT for the first time (group 3) were selected for further analysis as described in our experimental design earlier (Fig. 1). Each experimental condition was examined in three biological replicates, generating a total of 18 samples analyzed by deep RNA-seq.

Data were analyzed using the bioinformatic workflow described in figure S4. The produced reads were mapped against the *Leishmania mexicana* MHOM/GT/2001/U1103 genome of reference. Raw read counts per sample ranged from 25,521,644 to 32,949,665 reads. After QC, an average of 807,925 reads were discarded and the mean number of reads retained was 28,641,222. No sequences were flagged for poor quality by FastQC. Mean per-sequence and per-base accuracies were greater than 99.98% (Q > 38) before and after QC. Following alignment to the reference genome and gene-level quantification, an average of 23,918,544 (83.5%) of reads were assigned to annotated genes. After filtering out lowly expressed genes, an average of ∼8,353 genes were considered for each group comparison. The number of significant DEGs (FDR<0.05) for each comparison ranged from 2,136 to 5,419. These included 974 to 2,648 positively differentially expressed genes and 1,162 to 2,771 negatively differentially expressed genes per comparison.

To identify gene expression signatures associated with persistence we performed a bioinformatic analysis of high-throughput RNA sequencing based on five comparisons (Fig. 3).

**Figure 3.**
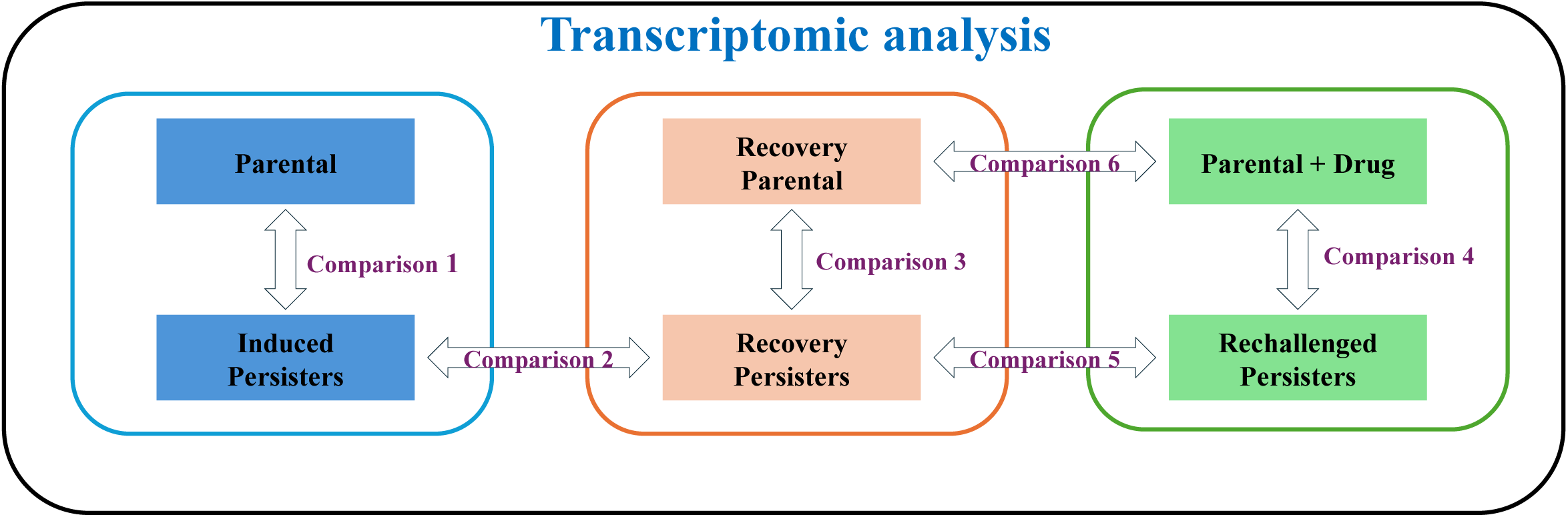
Bioinformatic comparison strategy for transcriptomic analysis. Schematic representation defining six specific differential gene expression comparisons used to dissect the molecular mechanisms of persistence in *Leishmania mexicana* deep RNA-seq samples. This analysis facilitates the characterization of distinct biological states across the induction, recovery, and re-challenge phases. Comparison 1 (Induction Response); Comparison 2 (Resuscitation in Persisters); Comparison 3 (Resuscitation Phase in Persisters vs parental control); Comparison 4 (Drug tolerance Memory); Comparison 5 (Drug response in persister parasites); Comparison 6 (Drug response in parental parasites).

Comparison 1 (Induced persisters vs Parental) investigates the initial state of surviving persister cells relative to the untreated parent population. This analysis enables the characterization of stress response pathways and treatment-induced dormant states, establishing a baseline comparison to identify the immediate transcriptomic signatures of survival and stress tolerance under initial PAT pressure during formation of persisters.

Comparison 2 (Recovery persisters vs Induced Persisters) allows us to understand the molecular mechanisms that drive exit from the persistent state following drug removal. This analysis of the reactivation dynamics shows specific transcriptomic shifts required to exit metabolic quiescence and resume active proliferation.

Comparison 3 (Recovery persisters vs Recovery Parental) identifies transcriptomic changes in persisters during recovery or resuscitation phase relative to control that was not exposed to drug (Recovery Parental). It characterizes the gene expression following drug withdrawal, assesses if persisters retain a molecular memory of prior treatment and evaluates whether surviving parasites remain different from the parental population once the stressor is removed and cells resume normal growth and increased metabolic activity.

Comparison 4 (Rechallenged persisters vs Parental + Drug) allows the determination of drug tolerance memory by investigating whether previously exposed parasites have a molecular memory of prior exposures and react differently to the drug pressure compared to a population facing it for the first time.

Comparison 5 (Rechallenged persisters vs Recovery Persisters) focuses on the molecular adaptations that occur when persister parasites after recovery phase are re-exposed to stressful, drug-induced conditions. This comparison is essential for identifying the specific gene expression shifts that enable enhanced tolerance and quiescence.

Comparison 6 (Parental + Drug vs Recovery Parental) evaluates the transcriptomic response of sensitive parental parasites to drug stress. By comparing treated and untreated populations, this analysis aims to identify the molecular mechanisms driving the response of sensitive parasites to the drug pressure. Further analysis of differences in drug responses in parental parasites and rechallenged persisters (comparison 5 vs 6) allows to determine the essential transcriptomic signatures associated with dormancy and drug tolerance.

Differential gene expression analysis revealed a reprogramming at the transcriptome level between persister-like and parental *Leishmania* populations across the evaluated conditions (Fig. 4). Number of genes reported herein were selected based on a FDR transformed p < 0.05. Bioinformatic analysis of immediate transcriptomic changes in parasites that survived initial drug treatment compare to untreated parental parasites (Comparison 1; Induced persisters vs Parental) revealed that a total of 3,124 genes were differentially expressed (Fig. 4). Of those, 1,537 genes were downregulated, suggesting the supression of different biological processes in order to survive the stress caused by the drug. It is typically associated with a strategy where parasites reduce their metabolic activity and proliferation to face drug exposure (27). Likewise, 1,587 genes were upregulated, which could be an indicator of activation of specific genes associated to drug stress response and involved in survival pathways, which are required for the establishment of the persistent population (28).

**Figure 4.**
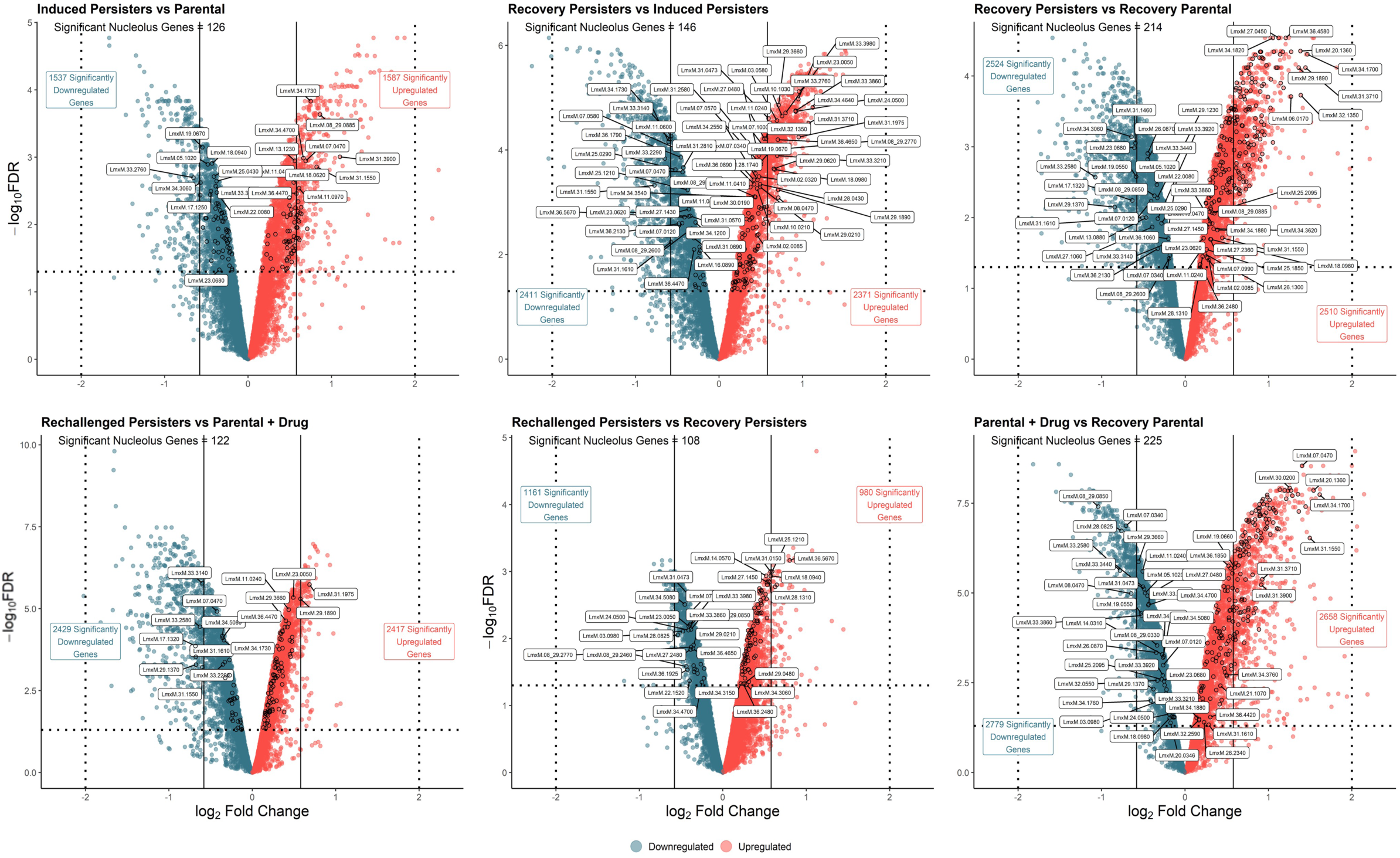
*Leishmania mexicana* persisters exhibit distinct gene expression profiles compared to parental parasites. Volcano plots showing the differential gene expression for the six specified experimental comparisons (Comparisons 1-6) based on statistical significance thresholds. The x-axis represents log2 fold change (log2FC), and the y-axis shows the statistical significance as the negative base-10 logarithm of the adjusted p-value (-log10 FDR). The horizontal dashed line indicates the statistical significance threshold (FDR < 0.05). Significantly upregulated genes are highlighted in red, while significantly downregulated genes are indicated in blue. Dots under the horizontal dashed threshold represent non-significant transcripts. Specific labeled data points correspond to key genes associated with the nucleolus. Each plot summarizes changes at the transcriptome level associated with the transition between parental and persisters-like states. Comparisons 1-6 in raw data are designated as C1-C6. The raw data are available in Source Data 2 (S2 Data).

The direct comparison between Recovery Persisters and Induced Persisters (Comparison 2) provides insights into how the parasite exits dormancy following drug stress. This contrast defines a “transcriptomic recovery” phase, characterized by the significant upregulation of 2,371 genes and the downregulation of 2,524 genes. Crucially, the upregulated transcriptomic profile offers a direct window into the specific molecular pathways that the parasite must reactivate to successfully exit latency and resume cellular proliferation. Examination of induced persisters in drug-free medium during recovery stage (Comparison 3; Recovery persisters vs Recovery Parental) revealed that even after 12 hours of recovery in drug-free medium transcriptome of persisters was drastically different with 2,510 genes upregulated and 2,524 downregulated compared to parental parasites that were never treated with drug. Analysis of rechallenged persisters and parental cells exposed to the drug first time (Comparison 4; Rechallenged persisters vs Parental + Drug) demonstrated that 2,417 genes were upregulated and 2,429 genes were downregulated in rechallenged persisters. This differential gene expression indicates that recovery persister parasites respond to drug rechallenge in a drastically different manner compared to the parental population facing the drug for the first time. Our findings suggest that prior drug exposure alters the persister population in such a way that key survival genes remain differentially expressed even after drug removal. This persistent gene expression acts as a “molecular memory” that equips persisters to better handle subsequent drug exposure compared to naive parental cells.

Analysis of drug response in persisters upon rechallenge compare to recovery persisters (Comparison 5; Rechallenged persisters vs Recovery Persisters) reveals that fewer genes were differentially expressed, (980 Up vs 1,161 Down) again suggesting the presence of a molecular memory in the parasites that allow them to be pre-adapted to withstand drug exposure. Due to this, the rechallenge does not trigger a large temodeling at the transcriptome level, because some key survival pathways could remain active. This provides additional support for the persistence of a molecular memory in the persister populations.

Comparison 6 (Parental + Drug vs Recovery Parental) analyzed the response of drug naïve parasites upon initial exposure. In contrast to Comparison 5, this comparison captures the immediate, acute response of drug-naïve parasites facing drug stress reflecting a primary stress response characterized by a high number of differentially expressed transcripts with 2,658 transcripts upregulated and 2,779 downregulated. When analysed alongside comparison 4, we observed drastic differences in transcriptomes between naïve and previously exposed populations.

Across comparisons, persister-like parasites displayed a substantial number of differentially expressed transcripts, indicating a response to drug-induced stress. Notably, both upregulated and downregulated gene sets were observed, suggesting that persistence is associated with coordinated activation and repression of distinct biological pathways rather than a unidirectional transcriptome shift. These results support a global remodeling of the transcriptome during the transition to the persister state.

Given the global remodelling of the transcriptomes observed across all the comparions, we next sought to determine whether specific functional gene categories were preferentially affected in persisters. Because the nucleolus is known to drive transcriptome remodeling by coordinating ribosome biogenesis and acting as a stress sensor (29) we analyzed our bioinformatic data for differential expression of nucleolar genes. Bioinformatic analysis revealed differential expression for many nucleolar genes across all tested conditions as shown on volcano plots, supporting altered ribosome biogenesis and importance of translational control in the formation of persisters (Fig. 4). This control may directly influence mRNA stability by enabling selective translation; during stress, specific stress-response transcripts evade degradation and are actively translated (30). Conversely, general transcripts are repressed and targeted for subsequent degradation through the interaction of RNA-binding proteins with their 3’ untranslated regions (31).

In many organisms, stress conditions promote selective translation through remodeling of ribosomal protein composition (32, 33). To further understand whether the persister state involves modulation of the translational machinery components, we focused on transcripts encoding ribosomal proteins (RPs) (Fig. 5). Differential expression analysis revealed a distinct and non-uniform pattern of RP transcript regulation in persister-like parasites compared to parental controls. Unlike the global gene expression trends, RP transcripts exhibited a more nuanced distribution, with specific subsets of paralogs being selectively upregulated while others were downregulated during induction phase (Induced persisters vs Parental). This heterogeneous pattern suggests a potential reconfiguration of ribosomal composition rather than a simple reduction or increase in ribosome biogenesis, which could be associated with the formation of specialized ribosomes that contribute to translational control under stress conditions. Further remodeling of protein synthesis machinery was observed in persisters upon drug removal, with 90 ribosomal protein transcripts upregulated and only 2 downregulated (Recovery persisters vs recovery parental). Interestingly, when persisters were rechallenged with the drug for second time, they exhibited rather a drastic downregulation (22 transcripts downregulated vs 2 transcripts upregulated) of protein synthesis machinery compared to the induction phase (Induced persisters vs Parental) and parental parasites (Parental + drug vs Recovery Parental). This downregulation correlated with the increase in the drug tolerance observed upon rechallenge (Fig. 2).

**Figure 5.**
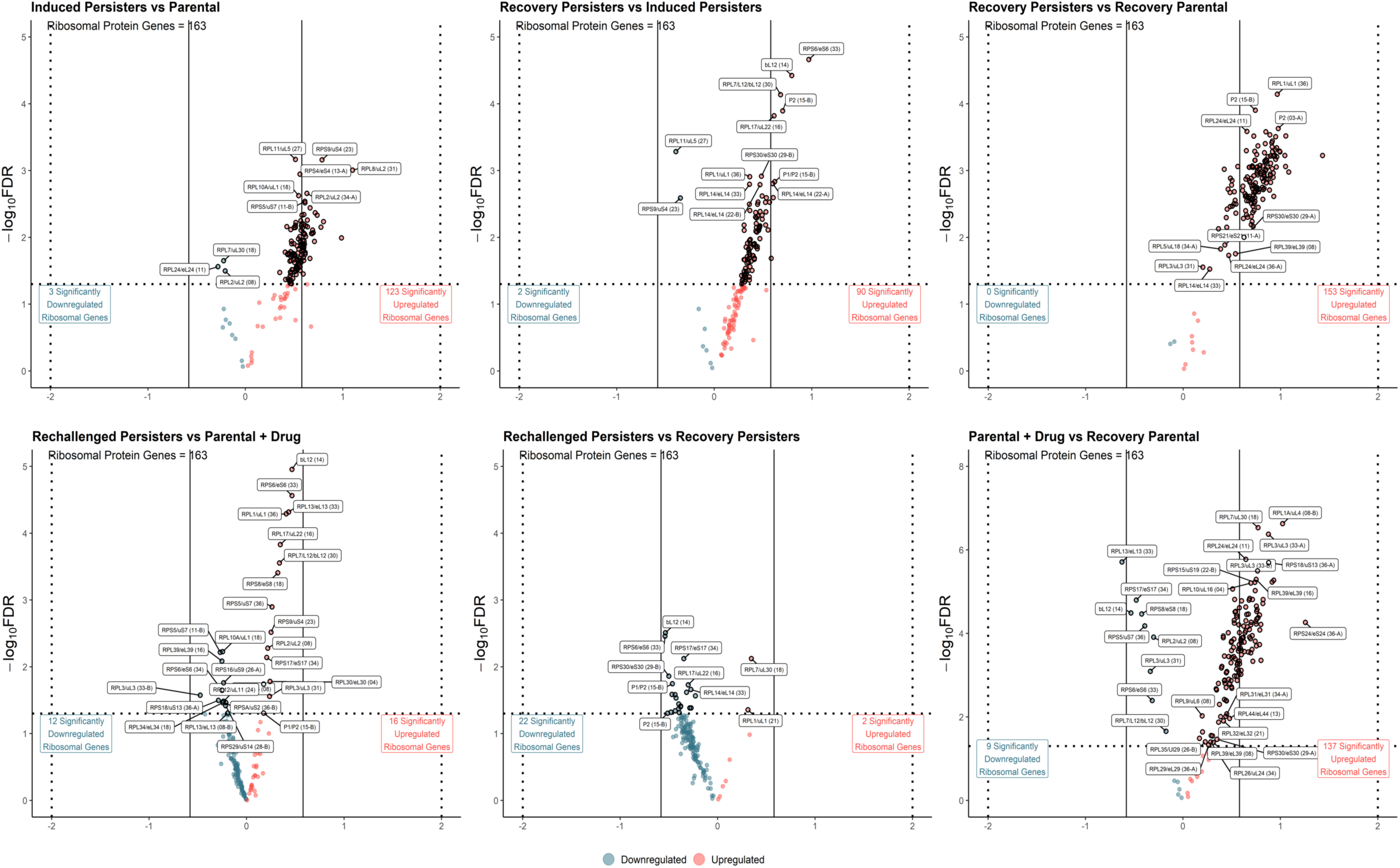
*Leishmania mexicana* persisters exhibit a different expression pattern of ribosomal proteins transcripts compared to parental parasites. Volcano plots showing differential expression of genes encoding ribosomal proteins (RPs) between the six evaluated comparisons (Comparisons 1-6). The x-axis represents log2 fold change (log2FC), and the y-axis shows the statistical significance as the negative base-10 logarithm of the adjusted p-value (-log10 FDR). The horizontal dashed line indicates the statistical significance threshold (FDR < 0.05). Significantly upregulated genes are highlighted in red, while significantly downregulated genes are indicated in blue. Dots under the horizontal dashed threshold represent non-significant transcripts. The distribution highlights selective regulation of specific RP paralogs, suggesting potential remodeling of ribosomal composition. 163 genes encoding RPs were evaluated. The raw data are available in Source Data 3 (S3 Data).

### Overrepresentation analysis uncovers multifaceted adaptations and unique pathway expression signatures in persisters

To determine coordinated functional trends and pathways in persisters, we performed Overrepresentation Analysis (ORA) across all the comparissons (Fig. 6). Normalized Enrichment Scores (NES) demonstrated consistent enrichment patterns across Biological Process (BP), Cellular Component (CC), and Molecular Function (MF) ontologies. The enrichment patterns revealed a consistent overrepresentation of pathways linked to ribosome biogenesis, rRNA processing, and nucleolar function, particularly in comparisons involving the induction and recovery phases. In general, the acute phases of drug response (Induced persisters and Parental + Drug) are characterized by a profound global shutdown of multiple biological processes. This widespread downregulation serves as a primary survival mechanism to withstand drug toxicity, being notably more pronounced in the induced persister population, which reflects a highly specific and coordinated stress response. Conversely, the transcriptomic landscape shifts during the recovery phases (Comparisons 2 and 3). Upon transfer to a drug-free medium, we observed a broad upregulation across numerous pathways such as an increase in ribosome production, cellular biosynthesis and mitochondrial function. This transcriptional reversal clearly captures the parasites exiting dormancy, representing an active biological effort to reactivate metabolism and resume normal growth dynamics. Meanwhile, comparisons evaluating the second drug response in rechallenged persisters (Rechallenged Persisters vs. Recovery Persisters, and Rechallenged Persisters vs. Parental + Drug) demonstrate a rapid return to metabolic quiescence. Upon re-exposure to the stressor, these parasites exhibited a severe downregulation of ribosomal pathways alongside a broad suppression of core regulatory networks, including general gene expression, macromolecule biosynthesis, and global metabolic processes. This indicates a primed, highly efficient repression of energy-intensive anabolic pathways to ensure survival during recurring stress. These findings are in agreement with the differential expression patterns observed for ribosomal proteins transcripts (Fig. 5), reinforcing the central role of translational regulation in the persister phenotype.

**Figure 6.**
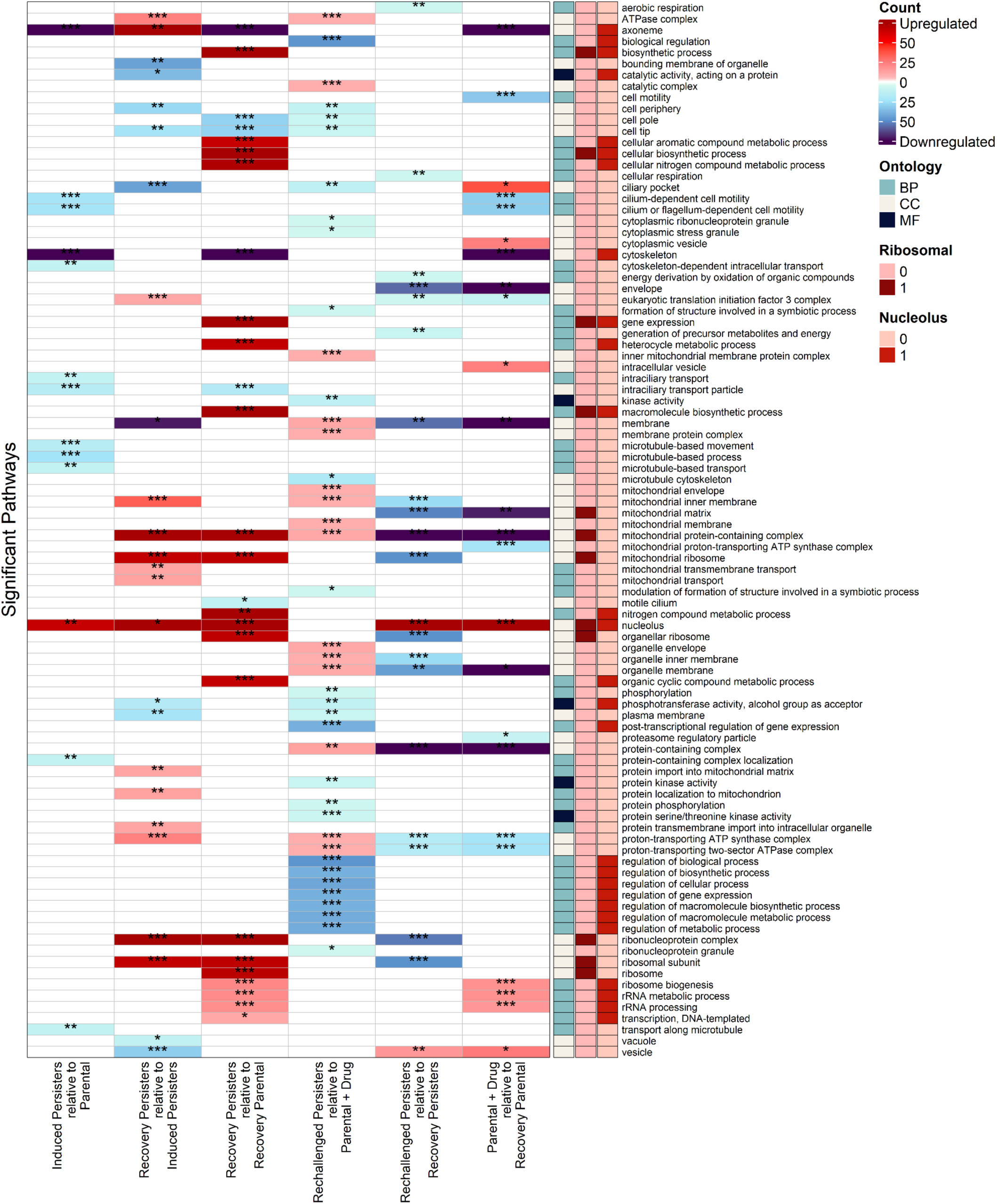
*Leishmania mexicana* persisters exhibit distinct pathways expression profiles compare to parental parasites. Overrepresentation Analysis (ORA) comparing all the evaluated groups. Cells show the Normalized Enrichment Score (NES) for Gene Ontology (GO) terms associated with Biological Processes (BP), Cellular Components (CC), and Molecular Functions (MF). Cells are differentially colored based on wheter they were significantly upregualted or down regulated among the evaluated groups. GO terms were annotated based on their association to ribosomal or nucleolus processes. Degree of significance within cells is denoted by the following p-value thresholds. * = p < 0.05; ** = p < 0.01; *** = p < 0.001. The raw data are available in Source Data 2 (S2 Data).

These results indicate that the transcriptome remodeling observed in persister-like parasites is functionally coherent and strongly biased toward processes involved in protein synthesis and ribosome dynamics to coordinate multifaceted changes in metabolism and other processes.

### Persisters exhibit distinct drug response profiles driven by the downregulation of protein synthesis machinery and upregulation of drug-response genes

To determine differences in drug responses of drug-tolerant persisters, we compared rechallenged persisters with drug-naïve parental parasites exposed to the treament for the first time (Comparison 4 vs Comparison 5, Fig. 7). The venn diagram (Fig. 7A) illustrates both shared and unique sets of differentialy expressed genes between conditions, highlighting that rechallenged persisters exhibit a distinct response. Our bioinformatic analysis revealed that 241 (197+44) transcripts are uniquely upregulated only in rechallenged persisters versus parental control treated with drug for the first time (Fig. 7A). Among them, 44 upregulated transcripts were downregulated in treated parental control. Similarly, rechallenged persisters exhibited 316 (255+61) uniquely downregulated transcripts, 61 of which were upregulated in control parasites upon drug exposure. Thus, we identified unique differences in drug responses in rechallenged persisters that could contribute to the increased drug tolerance. Interestingly, drug-naïve parental parasites responded to the drug in a broader way with 1,858 transcripts uniquely upregulated and 1,890 being downregulated. It is possible that due to the molecular memory in rechallenged persisters, many changes remain in place and therefore, driving to a coordinated and targeted response.

**Figure 7.**
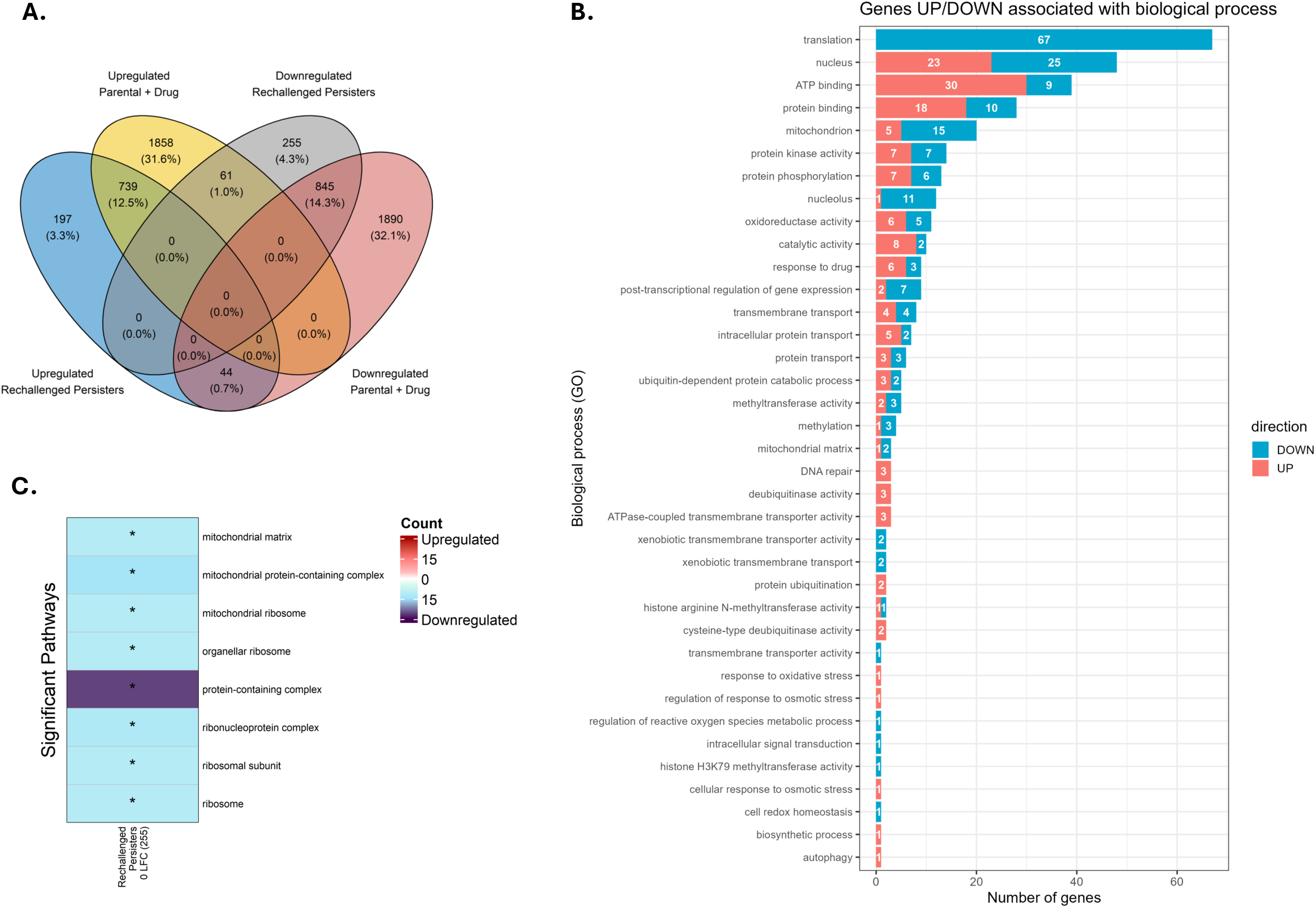
*Leishmania mexicana* persisters exhibit distinct drug response profiles compared to parental parasites. **(A)** Venn diagram comparing upregulated and downregulated genes among the groups ‘Parental + Drug’ and ‘Rechallenged persisters’. Percentages are calculated based on the total number of genes significantly upregulated and downregulated among the two groups (5,889). **(B)** Biological pathways associated with uniquely upregulated (197+44) or downregulated (255+61) genes in rechallenged persisters in response to drug. Hypothetical genes without known function were removed from the analysis (255 genes). The plot highlights major biological processes altered during the persister response. The translation term is the result of the combining of different GO terms, all of which are related to the translation machinery (GO:0006412, GO:0003743, GO:0006414, GO:0003735, GO:0005840, GO:0003723) **(C).** Biological processes enriched among uniquely downregulated genes in rechallenged persisters (255). Significant pathways associated with quiescence state were detected in this set of genes. The raw data are available in Source Data 4 (S4 Data).

Further analysis of these unique responses in rechallenged persisters demonstrated that these genes are associated with diverse biological processes involved in the parasite’s response to the drug pressure (Fig. 7B). Among these pathways, it is important to highlight the downregulation of the translation process and mitochondrial function, which is consistent with the previous results observed in the ORA analysis. Other relevant pathways included the enriched response to drugs and xenobiotic factors, as well as the response to oxidative and osmotic stresses.

The application of functional enrichment analysis to the uniquely downregulated genes (Fig. 7C) showed that these genes are represented in significant pathways associated with translation, ribosome and mithochondrial matrix, indicating the importance of translational control and repressing processes potentially linked to growth and proliferation, leading to dormancy in parasites and suggesting a specialized response to drug exposure.

Among the genes differentially expressed in rechallenged persisters in response to drug compared to first-time treated parental parasites, we prioritized those involved in drug response pathways due to their critical role in stress adaptation (Fig. 7A). We then manually curated this subset to identify genes that discriminate persisters from drug-sensitive parasites during response to the drug (Table 1).

**Table 1.** Genes associated with drug response in *Leishmania* persisters.

| Gene ID | Gene name | Direction | Rechallenged persisters<br>(Fold change) | Function | REF |
| --- | --- | --- | --- | --- | --- |
| LmxM.30.0020 | Aquaglyceroporin 1 (AQP1) | DOWN | 0.8296 | Drug uptake | (26) |
| LmxM.22.1420 | Aquaporin | DOWN | 0.8064 | Drug uptake | (26) |
| LmxM.24.2320 | Mitogen-activated protein kinase | DOWN | 0.8193 | Adaptive resistance | (34, 35) |
| LmxM.23.0250 | ABC-thiol transporter | UP | 1.1914 | Drug efflux | (26) |
| LmxM.30.1270 | p-glycoprotein e | UP | 1.1941 | Drug efflux | (35, 36) |
| LmxM.23.0220 | ABC transporter- Multidrug resistance protein | UP | 1.3197 | Drug efflux | (26) |

Interestingly, we found that genes associated with drug uptake, such as aquaglyceroporins (AQPs), were uniquely downregulated in persisters. In contrast, efflux pumps, including ATP binding cassette (ABC) transporters, were upregulated, presumably to remove the antimony drug from persister cells. These findings support the presence of a molecular memory in persisters-like parasites that allows a differentiated and efficient response to drug exposure.

## DISCUSSION

This study establishes a model to investigate the formation of persisters and the molecular mechanisms of persistence in *Leishmania mexicana*. Our findings demonstrate that persister-like parasites undergo coordinated, dynamic gene expression remodeling to promote drug tolerance, suppress proliferation, and activate adaptive stress responses.

By utilizing Ficoll gradient centrifugation following exposure to a lethal dose of potassium antimonyl tartrate (PAT) that kills 80% of parasites, we successfully isolated the viable subpopulation of persisters from the dead cells (27). Phenotypic characterization of this enriched fraction confirmed the hallmark traits of persistence: a significant growth delay (lag phase) upon transfer to drug-free medium and a markedly increased tolerance to subsequent PAT exposure (Fig. 2, S1_Data). Crucially, the ability of these parasites to eventually resume growth underscores the transient, non-heritable nature of this phenotype. Furthermore, these physiological adaptations are directly reflected in our bioinformatic analysis (Fig. 3).

To understand the molecular mechanisms driving the persister phenotype, we designed a comparative bioinformatic strategy encompassing six specific biological comparisons. This structured approach allowed us to isolate the transcriptomic signatures of distinct survival phases: the immediate stress response under initial drug pressure (Comparison 1), the exit of dormancy stage (Comparison 2), the metabolic signature governing the delayed phase during recovery (Comparison 3), the transcriptomic “memory” by evaluating whether previously exposed parasites have differences in transcriptomes upon a subsequent drug challenge compared to a naïve population (Comparison 4), the molecular adaptations that occur when persister parasites, after recovery phase, are re-exposed to stressful drug-induced conditions (Comparison 5), and finally, examination of the transcriptomic response of non-persisters, sensitive parental parasites exposed to drug stress (Comparison 6). During the acute stress phases (Comparisons 1 and 6), parasites entered a state of targeted metabolic quiescence, defined by a drastic suppression of many biological pathways coupled with nucleolar hyperactivity, probably needed to remodel transcriptome and the mobilization of primary defense effectors, such as heavy metal transporters (Fig. 4, S2_Data). Importantly, we demonstrated that this dormant state is fully reversible; upon drug removal (Comparisons 2 and 3), persisters exhibited a synchronous transcriptomic recovery, rapidly downregulating acute stress responses while reigniting ribosomal biogenesis to resume active proliferation. However, the true importance of this phenotype was revealed during the second drug challenge (Comparisons 4 and 5). Rechallenged persisters mounted a highly efficient, pre-adapted response—driven by epigenetic remodeling and catabolic reprogramming, that functionally phenocopies stable genetic resistance. Together, these stage-specific signatures demonstrate that *Leishmania* persistence is a highly plastic, actively regulated cycle of latency and transcriptomic memory, serving as a critical evolutionary bridge between transient survival and permanent drug resistance.

The extent of the transcriptomic remodeling is visually reflected in our global differential expression profiles. Volcano plots show the shifts in gene expression across all evaluated groups (Fig. 4). Detailed analysis of these shifts revealed the significant upregulation of several gene families classically associated with stress adaptation and drug survival. For instance, specific phosphatases (LmxM.18.0440, LmxM.13.1510, and LmxM.31.0640) were highly upregulated in induced persisters (S2_Data). In *Leishmania*, the overexpression of phosphatases is frequently observed under stress conditions and in drug-resistant strains, likely supporting the increased energy demands of drug efflux mediated by ABC transporters (37). Additionally, we observe the significant upregulation of key genes associated with the antimonial stress response, notably a putative zinc transporter 3 (LmxM.28.1930) and mitogen-activated protein kinase 2 (LmxM.13.0440) (34). The induction of these specific transporters and kinases highlights their critical role in mitigating heavy metal toxicity and orchestrating early survival signaling pathways. Additionally, we observed a pronounced upregulation of multiple amastin surface glycoproteins (LmxM.08.0840, LmxM.08.0850, LmxM.24.1270, and LmxM.30.4052), primarily in the “Induced Persisters” group. Amastins are critical for surface remodeling during environmental adaptation (38) and have also been found to be upregulated in parasites that were resistant to antimonials (34). Interestingly, a subset of these genes (LmxM.08.0750, LmxM.30.0452) remained upregulated in the “Rechallenged Persisters” (Comparison 5, S2_Data), suggesting that maintaining this surface modification might be a crucial defense mechanism when parasites re-encounter stressors. Conversely, during the drug-free recovery phase (Recovery Persisters vs. Recovery Parental), amastin expression was generally downregulated, reflecting a dynamic reversion to a baseline physiological state once the stress is removed. Notably, the induction of persistence was accompanied by the upregulation of specific heat shock proteins, such as HSP70 isoform (LmxM.28.2780). The accumulation of HSPs is a well-documented hallmark of cellular stress and has been directly linked to antimonial resistance in *Leishmania* (39)

In *Leishmania*, drug stress induced by antimonials is known to trigger severe oxidative stress response, which alters the expression of genes associated with oxidoreductase activity, such as Cytochrome b5 and Cytochrome P450 (40). Consistent with this, our analysis revealed an acute response to the initial drug stressor. We found that the LmxM.30.1170 (cytochrome b5 Heme/Steroid binding domain containing protein) and LmxM.29.3550 (cytochrome p450) were significantly upregulated in both the Induced persisters (Comparison 1, S2_Data) and the Parental + Drug group (Comparison 6, S2_Data). Additionally, other genes associated with oxidoreductase activity (LmxM.33.3330 and LmxM.34.2560) were upregulated in Comparison 6, while a distinct set of b5 associated genes (LmxM.09.1500 and LmxM.30.1190) were uniquely upregulated in Induced persisters. This transcriptional shift highlights the role of specific oxidoreductase activity in the initial metabolic response and drug detoxification.

Conversely, this acute activation of stress responses is coupled with a targeted metabolic shutdown. We observed a marked downregulation of other cytochrome b5-associated genes, such as LmxM.26.1430, across the induction (Comparison 1) and tolerance memory phases (Comparison 4), as well as LmxM.30.1200 in the Comparison 6 group (Fig 4, S2_Data). Since certain cytochrome b5 networks are intrinsically linked to energy-demanding anabolic pathways like lipid elongation (41), this repression likely represents a survival strategy where the parasite shuts down high-energy consumption processes to endure drug toxicity. Interestingly, during the recovery period (Comparison 3), many of the previously upregulated P450 and b5 genes returned to basal levels, indicating that their activation is a direct and exclusive response to the presence of the drug. Furthermore, these pathways remained generally downregulated in the rechallenged persisters, suggesting a process of transcriptomic adaptation where the parasites have consolidated a tolerant state that no longer relies on this acute oxidoreductase route. This metabolic efficiency is further evidenced by the dynamic regulation of drug transporters during the drug-free recovery phases (Comparisons 2 and 3). By shutting down processes that are no longer necessary, the parasite conserves vital resources. A prime example is the putative zinc transporter 3 (LmxM.28.1930); while this gene was highly upregulated during the acute drug responses (Comparisons 1 and 6), it was significantly downregulated in both recovery comparisons. This rapid reversion illustrates a highly coordinated transcriptomic mechanism that swiftly deactivates energy-intensive detoxification routes once the stressor is removed.

The analysis between Rechallenged Persisters vs Parental + Drug, unveils a transcriptomic memory in parasites previously exposed to PAT (S2_Data). A feature of this pre-adapted state is the upregulation of epigenetic modulators, specifically genes like histone-lysine N-methyltransferases (LmxM.20.0030 and LmxM.32.1790) which belongs to the family of histone methyltransferases, enzymes that are required for cell cycle regulation, maintenance of genome integrity in response to DNA damage, and initiation of replication (42, 43). The upregulation of this gene suggests that persisters deploy targeted histone post-translational modifications to face the drug presence through epigenetic modifications (44). Interestingly, this epigenetic remodeling appears functionally linked to the metabolic requirements of the quiescent state. Rechallenged persisters upregulated a distinct suite of genes driving nutrient scavenging and catabolism. This includes Cathepsin L-like proteases for protein degradation (LmxM.08.1060, LmxM.08.1070), a lysophospholipase for lipid recycling (LmxM.24.1840), and a glutamine aminotransferase for amino acid utilization (LmxM.32.1330). Previous studies underscore a regulatory axis between these processes, demonstrating that the genetic ablation of specific methyltransferases directly disrupts the expression profile of catabolic enzymes such as LmxM.08.1070 and LmxM.32.1330 (45). Thus, epigenetic mechanisms likely orchestrate the metabolic shift required to sustain parasite viability during periods of translational dormancy and extreme environmental stress.

Finally, to counteract the severe oxidative burst typically induced by antimonial drugs, rechallenged persisters mounted a rapid antioxidant response (S2_Data). This was evident by the targeted upregulation of key redox effectors, including a glutaredoxin-like protein (LmxM.27.0810) and thioredoxin (LmxM.01.0270), response that was unique for rechallenged persisters. Acting as non-enzymatic scavengers of free radicals, these proteins play a crucial role in preserving intracellular redox homeostasis, forming an essential defense layer that prevents lethal oxidative damage while the parasite remains in its non-replicative survival state (46, 47).

The results of our Overrepressentation Analysis (ORA) reveal that translational control is a fundamental driver of the persistent phenotype in *L. mexicana* (Fig. 6), highlighting a profound alteration in the pathways associated with the nucleolus and ribosomal proteins. This massive restructuring of the translation machinery during induction stage (Fig. 5, S3_Data) supports the need to remodel the transcriptome that is followed by the global reduction of protein synthesis machinery; additionally, our data support that downregulation of protein synthesis machinery is an essential adaptive strategy to slow growth down, allowing them to tolerate drug-induced stress. Interestingly, this translational and metabolic repression is not unique to protozoa but represents a universally conserved survival strategy in various domains of life (8, 48, 49).

Our transcriptomic data supports the hypothesis that *Leishmania* employs a convergent evolutionary strategy to face PAT pressure. By dynamically modulating the composition of the nucleolus, responsible for ribosome biogenesis, and subsequently altering the abundance of specific ribosomal subunits, persistent parasites are likely to minimize the drug’s toxic effects on active metabolism. At the same time, this translational pause allows them to conserve energy and reallocate resources exclusively towards basic cellular maintenance, by selectively and preferentially translating key genes involved in stress response and survival, thereby preserving their viability until the environmental stress subsides and they can resume proliferation (9, 11). Furthermore, this nucleolar enrichment could reflect a protective sequestration mechanism associated with the nucleolus detention pathway. During severe xenobiotic stress, eukaryotic cells frequently utilize the nucleolus to dynamically sequester essential cell cycle and metabolic proteins, safeguarding them from drug-induced damage or aggregation until the stress is removed (50, 51).

Analysis of drug response genes in rechallenged persisters identified several genes that were uniquely altered in response to antimonial drug, differentiating persisters from drug-naïve parasites and supporting presence of a regulated process required to survive drug pressure (Table 1, S4_Data). In *Leishmania*, the downregulation in the expression of membrane channels such as aquaglyceroporin 1 (AQP1, LmxM.30.0020) and aquaporin (LmxM.22.1420), alongside mitogen-activated protein kinases (MAPKs, LmxM.24.2320), is known to drive drug resistance by reducing cellular drug uptake (26). Likewise, resistance mediated by the overexpression of ABC transporters (LmxM.23.0250, LmxM.23.0220) and P-Glycoprotein (LmxM.30.1270), are key for drug efflux (26). The trend observed in the persister population suggests that these parasites might be using a similar coordinated transport mechanism to survive acute drug pressure.

Altogether, these findings demonstrate that drug persistence in *Leishmania* is a highly sophisticated, active survival strategy. This transient phenotype is orchestrated through precise transcriptome remodeling, coordinated translational downregulation, and distinct epigenetic adaptations. Moreover, this state likely represents the initial evolutionary steppingstone toward the acquisition of permanent genetic drug resistance, as previously described in bacterial models (52). By establishing the model to study persistence in this protozoan parasite, we could define the survival mechanism of *Leishmania* when facing drug pressure, while also highlight novel molecular vulnerabilities within the persister population, offering promising therapeutic targets to intercept the transition to resistance and ultimately prevent clinical relapse.

## Materials and Methods

### Parasite culturing

Promastigotes of *Leishmania mexicana* (MNYC/BZ/62/M379 GFP) were cultured at 26°C in complete M199 medium (M199 medium at pH 7.4 supplemented with 10% fetal bovine serum, 1.82 g/L sodium bicarbonate (NaHCO3), HEPES 40 mM (pH 7.4), 0.1 mM adenine, 1 mg/L biotin, 5 mg/L hemin, 2 mg/L biopterin 100 units/mL of penicillin and 100 μg/mL of streptomycin) (17). Parasites were seeded at 5×10^5^ cells/mL and incubated for 48 hours until reaching early log phase. Promastigotes in this phase were harvested and seeded to required concentration for each experiment henceforth.

### Estimation of EC_50_ and EC_80_ doses

Early log phase promastigotes were harvested and seeded at 2×10^8^ cells/mL in Schneider’s insect medium (Sigma, Cat: S0146) containing 10% FBS (Corning, Cat: 35-011-CV) and 1% Penicilin-Streptomycin (Sigma-Aldrich, Cat: P0781). Half maximal effective concentration (EC_50_) of parasites treated with potassium antimony (III) tartrate (PAT, Sigma-Aldrich, Cat: 230057) was estimated by MTT assay in 96-well plates under two experimental conditions: control samples with 50 µL of parasite culture combined with 50 µL of Schneider’s medium without drug, and treated samples including 50 µL of parasite culture combined with 50 µL of Schneider’s medium with serial dilutions of PAT (2, 1, 0.5, 0.25, 0.125 and 0 mM). Schneider’s medium without drug was used as blank. Three biological repeats each with three technical repeats were analyzed. Plate was incubated at 26°C for 24 hours, after drug challenge, 20 µL of MTT solution at 5 mg/mL (Sigma-Aldrich, Cat: M2128) were added to each well to evaluate parasite viability. After four hours treatment with MTT, reaction was stopped by adding 100 µL of 10% SDS-HCl solution and absorbance was measured at 570 nm. EC_50_ value was calculated by non-linear regression analysis on dose-response curve using GraphPad Prism 9. EC_80_ was estimated with EC_50_ value and the slope factor using online GraphPad calculator (https://www.graphpad.com/quickcalcs/ECanything1/).

### EC_80_ Drug treatment

After 48 hours incubation under previously described conditions, early log phase culture was harvested and seeded at 2×10^8^ cells/mL in complete M199 medium with or without PAT according to the group: Control group included parasites without drug, while treated group was composed of parasites challenged with PAT at 0.315 mM, which is the concentration that kills 80% of parasite population (EC_80_). The cultures were incubated at 26°C for 24 hours and each group was analyzed in triplicates.

### Live Parasite Separation on Ficoll Gradients

Once the treatment finished, *Leishmania* cells with good viability were enriched with Ficoll gradients. Briefly, promastigotes were harvested and centrifuged at 3,000× *g* for 10 min. The pellet was resuspended in 4 mL of PBS. The suspension of cells was added as the last layer on a Ficoll gradient (Ficoll Type 400) prepared in 15 mL tubes containing 2 mL of Ficoll 20% at the bottom and 2 mL of Ficoll 10% on top. Gradients were centrifuged at 1,300× *g* for 15 min at room temperature. After centrifugation, the fraction of Ficoll 10% containing live cells was recovered and washed with 10 mL of M199 medium. After centrifugation at 2,500× *g* for 5 min, the pellet was resuspended in 1 mL of medium M199. Cell concentration was calculated, 2×10^7^ cells were centrifuged, and the pellet was resuspended in TRIzol and stored at -80°C until RNA purification. An aliquot of each replica was taken to check proliferation; these aliquots were considered as T0. An additional aliquot of each replica was used to examine resuscitation for 12 hours in drug free medium followed by cell collection for RNA extraction.

### Determination of persister’s tolerance to drug

To test the persisteŕs tolerance to different drug concentrations, an aliquot was taken after 12 hours of recovery. The number of cells was estimated, and parasites were seeded in 96-well plates at 5×10^6^ motile cells/mL in Schneider’s insect medium containing different known concentrations of PAT (2, 1, 0.5, 0.25, 0.125 mM) by triplicate. Plate was incubated for 24 hours at 26°C, then 20 µL of MTT 5 mg/mL were added per well to evaluate the parasite viability. After four hours of MTT treatment, the reaction was stopped by adding 100 µL of 10% SDS-HCl solution and absorbance was read at 570 nm. EC_50_ values were estimated using non-linear regression.

### Parasites proliferation after drug treatment

After EC80 drug treatment live parasites enriched on Ficoll gradients were collected and aliquot of each replica corresponding to *t=*0, including both control and treatment, was centrifuged at 2,500 x *g* for 5 minutes and washed twice with M199 medium to remove all drug remnant. Then, each culture was seeded at 2.5×10^5^ cells/mL in fresh M199 media without any drug for parasites recovery. Cultures were incubated at 26°C for 96 hours. Growth rate was monitored during parasite recovery by taking aliquots from each sample and counting by hematocytometer every 12 hours.

### RNA purification

RNA was isolated using a modified protocol from Invitrogen TRIzol LS protocol. 0.75 mL of TRIzol LS reagent was added per 0.25 mL of sample. Samples were homogenized by pipetting and incubated for 5 min to ensure a complete lysis. 0.2 mL of chloroform were added per 0.75 mL of TRIzol LS, the samples were incubated 3 min and centrifuged for 15 minutes at 12,000 × *g* at 4°C. After that the aqueous phase was recovered into a new tube and 1 µL of glycogen and 0.5 mL of isopropanol per 0.75 mL of TRIzol were added. Samples were incubated overnight at -20°C, washed with ethanol and dried RNA pellet was resuspended in 30 µL of nuclease-free water. Each sample was quantified by NanoDrop One (Thermo Scientific^TM^).

### High-throughput sequencing

Eighteen samples were processed for high-throughput sequencing of RNA (Deep RNA-Seq). Isolated RNA sample quality was assessed by High Sensitivity RNA Tapestation (Agilent Technologies Inc., California, USA) and quantified by AccuBlue® Broad Range RNA Quantitation assay (Biotium, California, USA). Paramagnetic beads coupled with oligo d(T)25 are combined with total RNA to isolate poly(A)+ transcripts based on NEBNext® Poly(A) mRNA Magnetic Isolation Module manual (New England BioLabs Inc., Massachusetts, USA). Prior to first strand synthesis, samples are randomly primed (5’ d(N6) 3’ [N=A, C, G, T]) and fragmented based on manufacturer’s recommendations. The first strand is synthesized with the Protoscript II Reverse Transcriptase with a longer extension period, approximately 40 minutes at 42⁰C. All remaining steps for library construction were used according to the NEBNext® Ultra™ II Non-Directional RNA Library Prep Kit for Illumina® (New England BioLabs Inc., Massachusetts, USA). Final libraries quantity was assessed by Qubit 2.0 (ThermoFisher, Massachusetts, USA) and quality was assessed by TapeStation D1000 ScreenTape (Agilent Technologies Inc., California, USA). Final library size was about 430bp with an insert size of about 300bp. Illumina® 8-nt dual-indices were used. Equimolar pooling of libraries was performed based on QC values and sequenced on an Illumina® NovaseqX plus platform (Illumina, California, USA) with a read length configuration of 150 PE for 40M PE reads per sample (20M in each direction).

### Data preparation for RNA-seq Analysis

FASTA, GFF, and GAF files for *Leishmania mexicana* reference strain MHOM/GT/2001/U1103 were downloaded from TriTrypDB v68 (53). The GFF file was converted to GTF, using the tool agat_convert_sp_gff2gtf.pl (54) to meet downstream requirements. fastp v1.0.1 (55) was used to process the paired raw FASTQ files, and arguments --trim_poly_g and --trim_poly_x were explicitly specified to trim polyG sequencing artifacts and polyA mRNA tails. FastQC v0.11.8 (56) was used to assess quality pre and post processing. Reports were aggregated using MulitQC v1.29 (57) STAR v2.5.2b (58) and used to index the reference genome and align clean reads. Aligned reads were subsequently quantified using featureCounts v2.1.1 (59). A custom *L. mexicana* annotation database was built using makeOrgPackage from the AnnotationForge package (60) in R v4.2.1 based on curated Gene Ontology (Biological Process, Cellular Component, and Molecular Function) annotations.

### Differential Expression Analysis and Functional Enrichment Analysis

Differential expression was assessed using the edgeR (v3.38.4) quasi-likelihood analysis pipeline (61) in R v4.2.1. Lowly expressed genes were excluded using filterByExpr (61). Library sizes were normalized with the trimmed mean of M values (TMM) method using calcNormFactors (61). Gene-wise dispersions were estimated with estimateDisp(), and quasi-likelihood negative binomial generalized linear models were fitted with glmQLFit()(61)Differential expression testing was performed using glmQLFTest (61) and p-values were adjusted for multiple testing using the Benjamini–Hochberg false discovery rate (FDR) procedure. Functional gene enrichment was conducted using clusterProfiler v4.4.4 (62) in R v4.2.1. Significant genes as defined by FDR<0.05 and were selected for overrepresentation analysis (ORA) using enrichGO (62). ORA was performed separately for positively and negatively differentially expressed genes. The set of genes considered for differential expression testing was used as the background gene list.

### Statistical analyses

Statistical analyses of the results were performed using GraphPad Prism v9.0, while analyses of RNA-seq results were done using R v4.2.1. Graphical visualization was performed using various R packages. “ComplexHeatmap” was used to visualize functional pathway analyses with modifications for cell annotation of p-values(63). For Venn diagram creation the R package “ggVennDiagram” was used(64)

## Supplemantry figure legends

**Figure S1.**
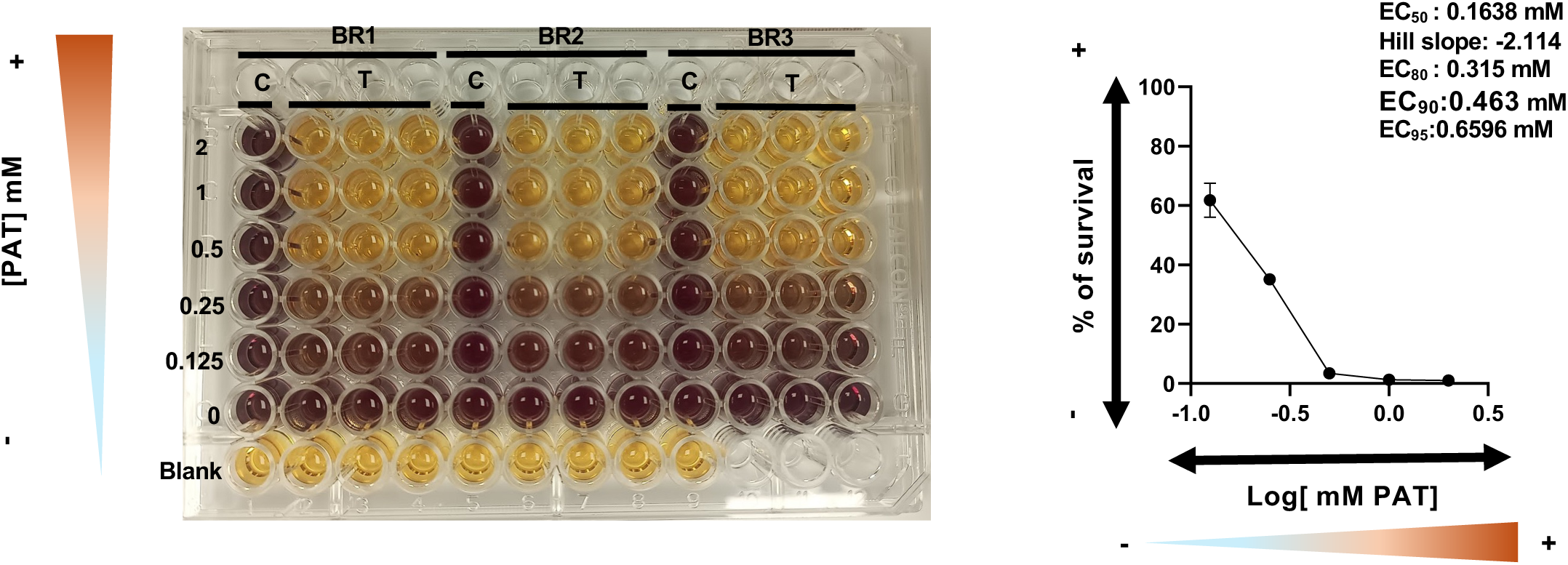
Half maximal effective concentration (EC_50_) for *L. mexicana* promastigotes treated with trivalent antimony. Effective concentration is the concentration (or dose) effective in producing a certain percentage of the maximal response and can be used to estimate the drug tolerance, **(A)** The colorimetric MTT assay was used to estimate the EC_50_ value. Three conditions were evaluated per experiment: parasites growing without SbIII (control, C), parasites growing under SbIII pressure (treatment, T), and medium without parasites (blank). The change from yellow (MTT) to purple (Formazan or reduced MTT) indicates cell viability. **(B)** The EC_50_ value was estimated by using a non-linear regression model (Sigmoidal model with four parameters). The X-axis is the log transform drug concentration. Y-axis represents the percentage of survival parasites and the arbitrary units of formazan obtained for the untreated parasites corresponded to 100%. Then the EC_80_ value was estimated based on the EC50 and Hill slope value by using the online calculator: https://www.graphpad.com/quickcalcs/ECanything2/. Trivalent antimony was administrated as potassium antimonyl tartrate (PAT). Three technical replicates and three biological replicates were performed (BR1-3). Mean and standard error are plotted. Degrees of Freedom: 13. R squared: 0.9711. Sum of Squares :244.3. Sy.x: 4.271. EC_50_ : 0.1638 mM. Hill slope: -2.114. EC_80_ : 0.315 mM. Lower (-). Higher (+).

**Figure S2.**
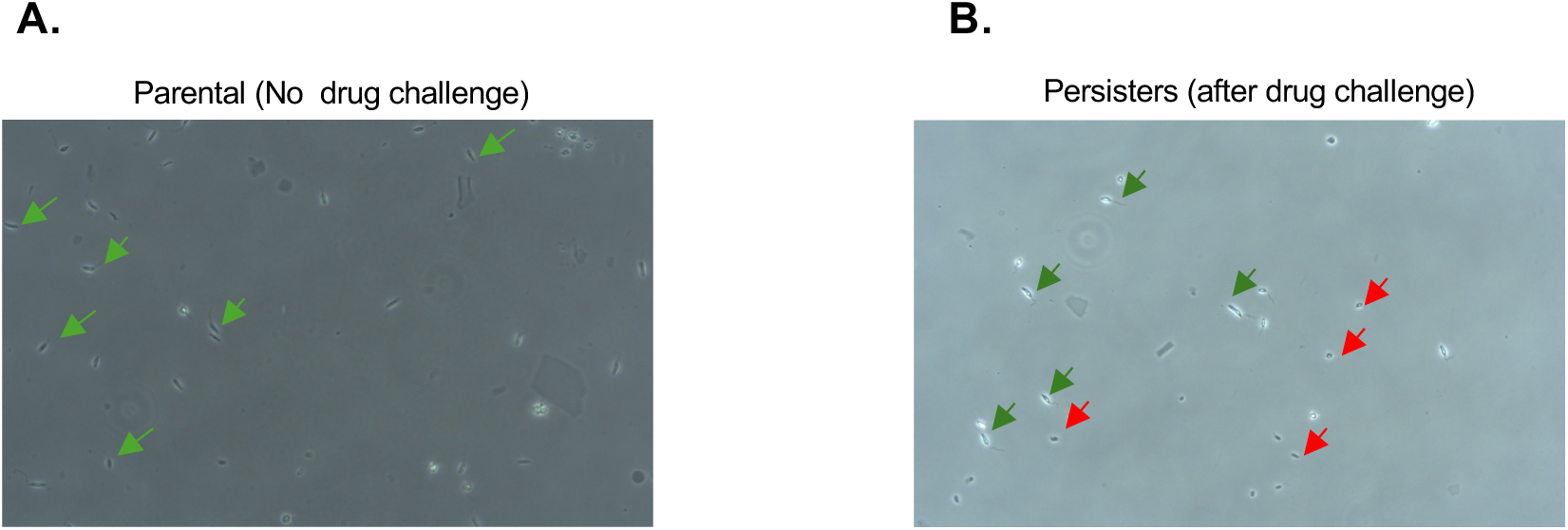
Persister parasites were identified after the drug challenge. Parental and persistent parasites were observed by using an inverted light microscope after the recovery stage (see figure1). (**A)** Parental or control group not treated with the drug showing normal parasite morphology (green arrows). (**B)** Drug treated group showing two different morphologies. Multiple apoptotic parasites (red arrow). A percentage of survival parasites with normal morphology and considered persisters (green arrow). Trivalent antimony was administrated as potassium antimonyl tartrate (PAT) at the EC_80_ value.

**Figure S3.**
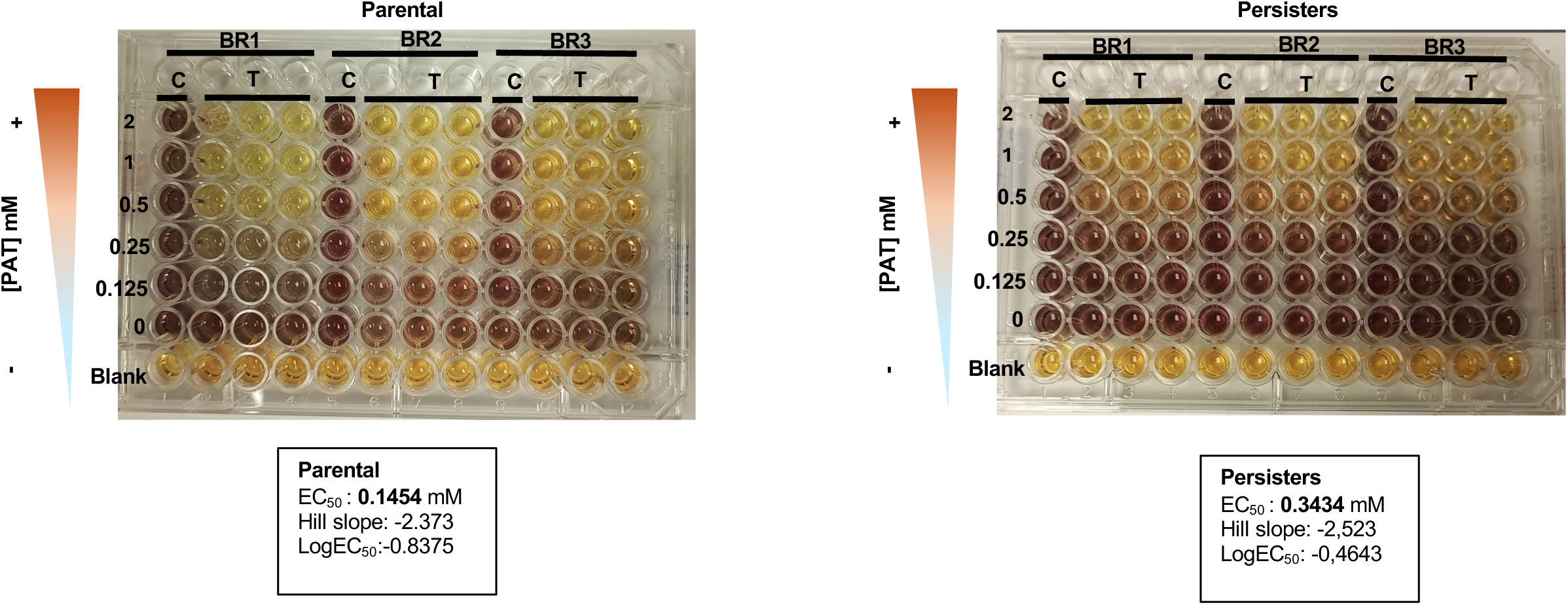
Half maximal effective concentration (EC_50_) for *L. mexicana* promastigotes treated with trivalent antimony after recovery with ficoll. Effective concentration is the concentration (or dose) effective in producing a certain percentage of the maximal response and can be used to estimate the drug tolerance**, (A)** The colorimetric MTT assay was used to estimate the EC_50_ value. Three conditions were evaluated per experiment: parasites growing without SbIII (control, C), parasites growing under SbIII pressure (treatment, T), and medium without parasites (blank). The change from yellow (MTT) to purple (Formazan or reduced MTT) indicates cell viability. (**B)** The EC_50_ value was estimated by using a non-linear regression model (Sigmoidal model with four parameters). The X-axis is the log transform drug concentration. Y-axis represents the percentage of survival parasites and the arbitrary units of formazan obtained for the untreated parasites corresponded to 100%. Then the EC_80_ value was estimated based on the EC50 and Hill slope value by using the online calculator: https://www.graphpad.com/quickcalcs/ECanything2/. Trivalent antimony was administrated as potassium antimonyl tartrate (PAT). Three technical replicates and three biological replicates were performed (BR1-3). Mean and standard error are plotted.

**Figure S4.**
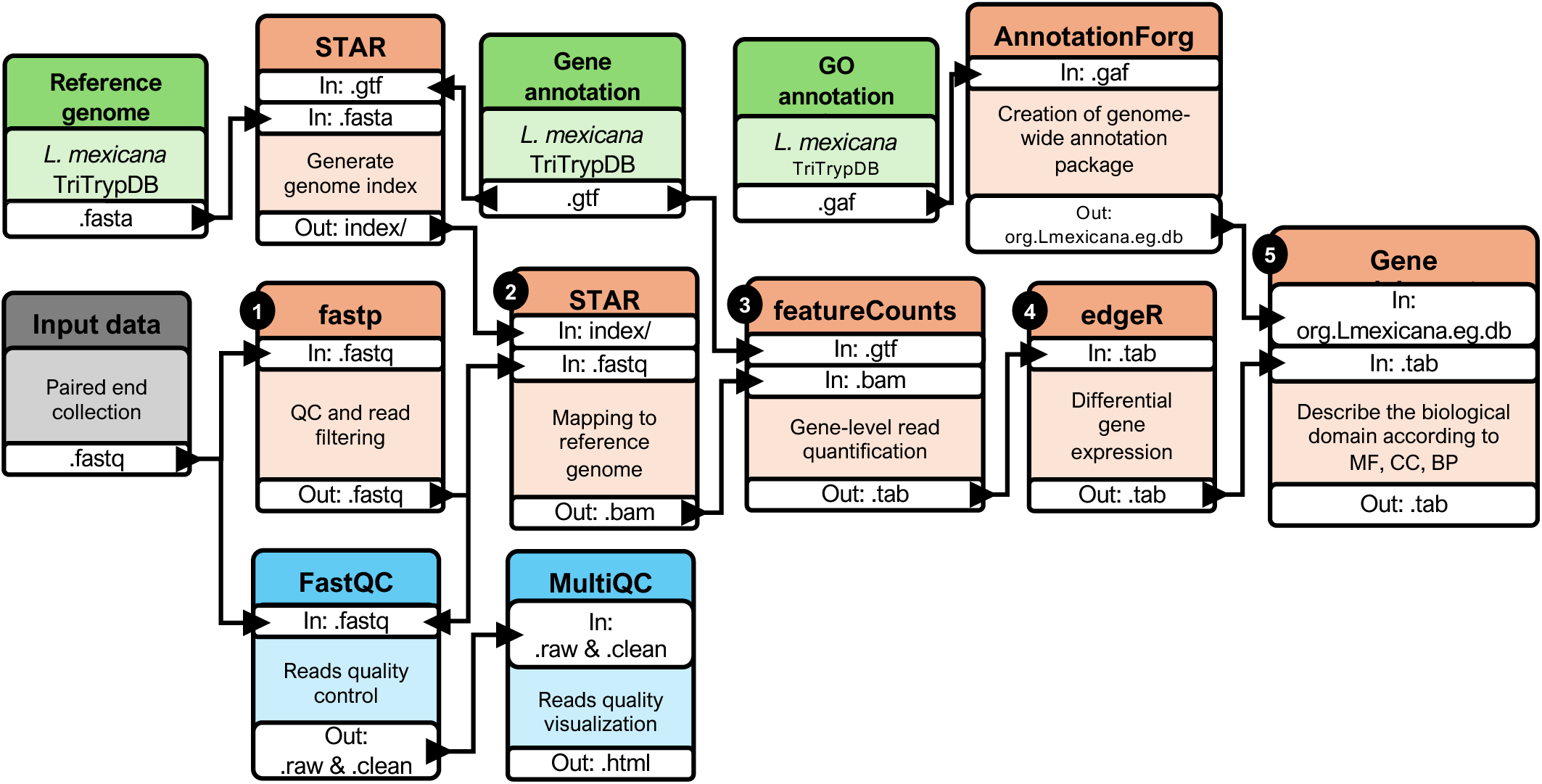
Bioinformatic workflow for RNA-seq analysis of Leishmania mexicana. Numbered boxes indicate workflow steps in logical order. Green boxes denote external reference files obtained from TriTrypDB and used during analysis. Gray box indicates raw input sequencing data. Tool names are shown in the box headers, the purpose of each step is described within the boxes, and the corresponding input (In) and output (Out) file types are indicated in the white panels. Yellow boxes denote read quality control assessment.

## Author Contributions

Conceptualization, Z.N.K. and E.Q.; methodology, E.Q. and Z.N.K.; data analysis, E.Q., C.C.R-A., K.O., J.A., C.P., Z.N.K.; resources, Z.N.K. and C.P.; writing - original draft preparation, E.Q., and Z.N.K.; writing—review and editing, E.Q., C.C.R-A, K.O., J.A., C.P., Z.N.K.; supervision, Z.N.K. and C.P.; project administration, Z.N.K and C.P.; funding acquisition - Z.N.K. and C.P. All authors have read and agreed to the published version of the manuscript.

## Funding

This study was funded by Institute for One Health Innovation seed grant (Texas Tech University; to Z.N.K. and C.P.) and Texas Tech University Health Science Center start-up funds (Z.N.K.).

## Data Availability Statement

The raw and processed data supporting the conclusions of this article are available in the Gene Expression Omnibus database from the NCBI under accession number GSE332713

## Acknowledgments

We thank members of the Karamysheva and Phillips laboratories for their advice and critical discussions. Deep RNA-seq was performed by Admera Health.

Part of figures 1 and 3 were created in BioRender.com.

## Conflicts of Interest

The authors declare no conflicts of interest.

## Supporting information

S1 Data. Source data for Figure 2. Raw values for growth curves and MTT assays.

S2 Data. Source data for Figure 4 and 6. Differential gene expression of comparisons 1-6 (C1-6).

S3 Data. Source data for Figure 5. Differential gene expression of ribosomal proteins for comparisons 1-6 (C1-6).

S4 Data. Source data for Figure 7. List of genes for venn diagram.

## Notes

### Competing Interest Statement

The authors have declared no competing interest.

